# Cxcl10 is required for survival during SARS-CoV-2 infection in mice

**DOI:** 10.1101/2024.09.30.613319

**Authors:** Shamik Majumdar, Joseph D. Weaver, Sergio M. Pontejo, Mahnaz Minai, Xinping Lu, Ji-Liang Gao, Gibran Holmes, Reed Johnson, Hongwei Zhang, Brian L. Kelsall, Joshua M. Farber, Derron A. Alves, Philip M. Murphy

**Author notes:** Corresponding author: Philip M. Murphy, M. D., Bldg 10, Room 11N111, NIH, Bethesda, MD 20892; (PMM).

## Abstract

Severe acute respiratory syndrome coronavirus 2 (SARS-CoV-2), the etiological agent of the coronavirus disease 2019 (COVID-19) pandemic, remains endemic worldwide ∼5 years since the first documented case. Severe COVID-19 is widely considered to be caused by a dysregulated immune response to SARS-CoV-2 within the respiratory tract. Circulating levels of the chemokine CXCL10 are strongly positively associated with poor outcome; however, its precise role in pathogenesis and its suitability as a therapeutic target have remained undefined. Here, we challenged 4-6 month old C57BL/6 mice genetically deficient in *Cxcl10* with a mouse-adapted strain of SARS-CoV-2. Infected male, but not female, *Cxcl10*^-/-^ mice displayed increased mortality compared to wild type controls. Histopathological damage, inflammatory gene induction and virus load in the lungs of male mice 4 days post infection and before death were not broadly influenced by Cxcl10 deficiency. However, accumulation of B cells and both CD4^+^ and CD8^+^ T cells in the lung parenchyma of infected mice was reduced in the absence of Cxcl10. Thus, during acute SARS-CoV-2 infection, Cxcl10 regulates lymphocyte infiltration in the lung and confers protection against mortality. Our preclinical model results do not support targeting CXCL10 therapeutically in severe COVID-19.

## Introduction

Since December 2019, the coronavirus disease 2019 (COVID-19) pandemic has caused over 7 million documented deaths, and is currently endemic worldwide, with almost 250,000 reported cases and more than 4,400 documented deaths in the last 28 days as of September 18, 2024 (https://data.who.int/dashboards/covid19/). Infection with the causative agent of COVID-19, severe acute respiratory syndrome coronavirus 2 (SARS-CoV-2), results in a broad spectrum of clinical manifestations ranging from asymptomatic infection to mild symptoms, long COVID or death in severe cases of acute respiratory distress syndrome (ARDS). Cytokine storm is cited as a major pathogenic factor for the development of severe COVID-19, consisting of heightened levels of inflammatory cytokines and chemokines, and the presence of inflammatory infiltrates, leading to ARDS and fatality ^1,2^.

The chemokine CXCL10 (C-X-C motif chemokine 10), previously known as IFN-γ– induced protein 10 (IP-10), binds to the receptor CXCR3, which is expressed on diverse cell types including naïve CD8^+^ T cells, recently activated naïve T cells, effector/memory subsets of CD4^+^ and CD8^+^ T cells, some γ/δ T cells and MAIT cells, NK cells, subsets of B cells, and plasmacytoid dendritic cells (https://www.proteinatlas.org/ENSG00000186810-CXCR3/immune+cell). CXCL10 levels in plasma and serum have been positively associated with the severity and progression of COVID-19 ^3–10^. Consequently, CXCL10 has been proposed as a prognostic marker and a potential mediator of COVID-19-induced ARDS. Therefore, targeting the CXCR3-CXCL10 axis is postulated to be therapeutic for patients with symptomatic COVID-19 ^11^. The aim of the present study was to investigate the direct role of CXCL10 during pathogenesis in mice infected with a mouse-adapted strain of SARS-CoV-2.

## Results

### Complete *Cxcl10 deficiency is a genetic risk factor for fatal outcome in male C57BL/6 mice infected with MA10 SARS-CoV-2*

To investigate the direct role of Cxcl10 during SARS-CoV-2 pathogenesis, we first infected *Cxcl10*^-/-^ C57BL/6 mice with the mouse-adapted 10 (MA10) variant of SARS-CoV-2, which causes acute morbidity and mortality in wild type mice ^12^. Mice were intranasally infected with an approximate LD_50_ dose of 8.62 × 10^4^ TCID_50_ of MA10 and observed for morbidity and mortality for a period of at least 12 days. In pilot studies, *Cxcl10*^-/-^ male mice showed increased mortality when compared to non-littermate JAX664 C57BL/6 mice as controls (Fig. 1A). Although the knockout and control mice are both on the C57BL/6 background, it is important to note the *Cxcl10*^+/+^ JAX664 C57BL/6 control mice are naturally genetically deficient in *Cxcl11*, a closely related chemokine that also signals through CXCR3, due to mutations in the *Cxcl11* gene that result in frameshift and truncated Cxcl11 protein ^13^. However, the *Cxcl10*^-/-^ mice on this background have the wild type (WT) *Cxcl11* gene due to homologous recombination of the *Cxcl10* knockout allele and the adjacent WT *Cxcl11* with genomic DNA from 129S4/SvLaeJ mice, which were used for construction prior to backcrossing onto the C57BL/6 line. Therefore, to control for Cxcl11 status and other unknown background genetic and microbiota differences, we infected littermate male *Cxcl10*^*-*/-^*Cxcl11*^+/+^ mice and *Cxcl10*^+/+^*Cxcl11*^+/+^ WT mice on the JAX664 background (see methods). Again, we observed higher mortality in the *Cxcl10*^*-*/-^ male mice than in WT control mice (Fig. 1B). In contrast, there was no difference in mortality in infected female *Cxcl10*^*-*/-^ *Cxcl11*^+/+^ mice compared with non-littermate *Cxcl10*^+/+^*Cxcl11*^-/-^ JAX664 C57BL/6 mice (Fig. 1C) or with WT *Cxcl10*^+/+^*Cxcl11*^+/+^ littermates (Fig. 1D). We also monitored the body weights of the mice during these survival studies. Both male and female, survivor and non-survivor *Cxcl10*^-/-^ mice, lost similar percentages of their initial body weights compared to the WT counterparts during the progression of infection (Fig. 1E-H). Therefore, complete genetic *Cxcl10* deficiency appears to be a risk factor for fatal outcome in male but not female C57BL/6 mice after acute MA10 SARS-CoV-2 infection. The subsequent experiments were all performed using male *Cxcl10*^+/+^*Cxcl11*^+/+^ (*Cxcl10*^+/+^) and *Cxcl10*^*-*/-^*Cxcl11*^+/+^ (*Cxcl10*^-/-^) littermates.

**Figure 1.**
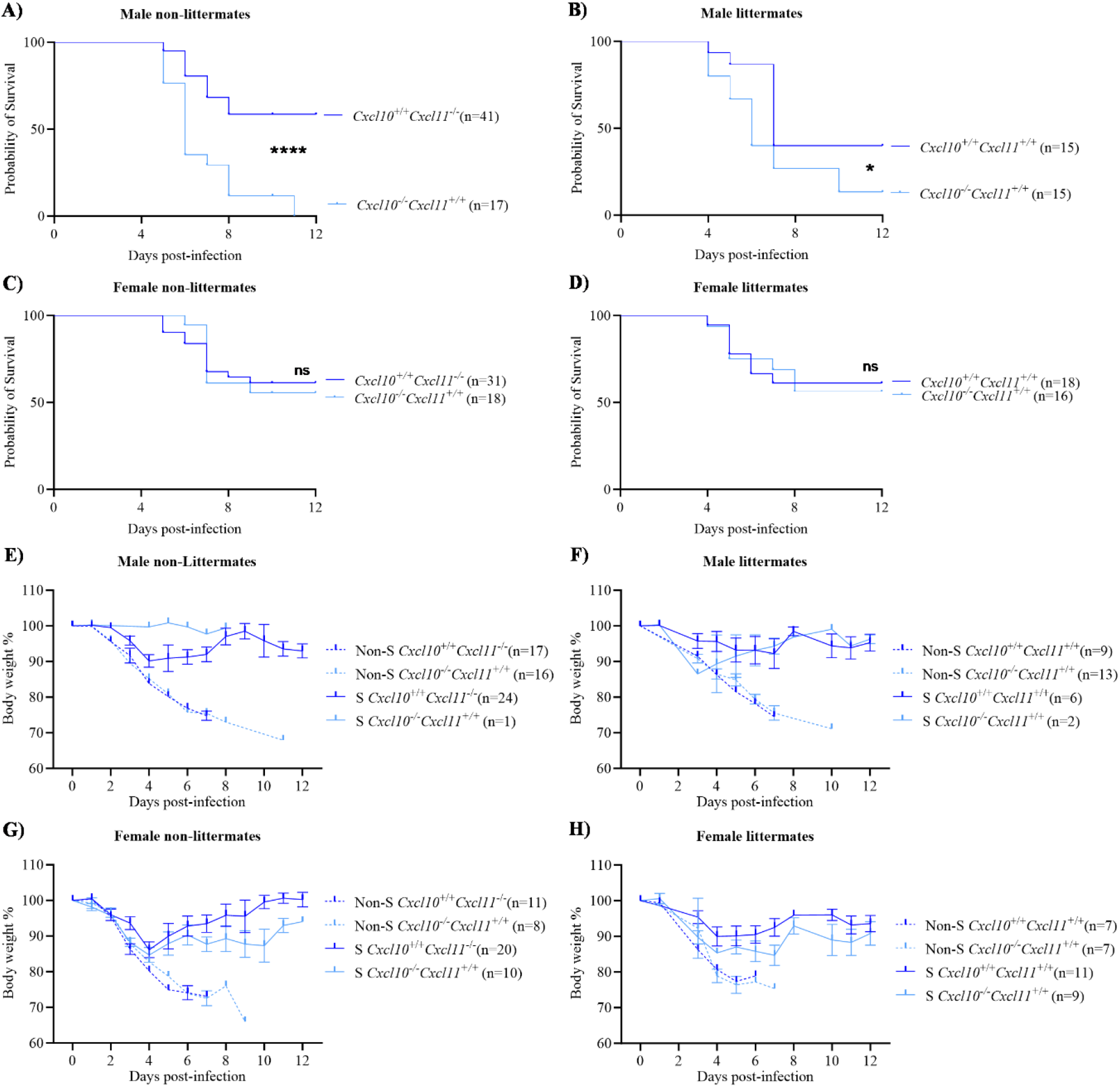
Complete Cxcl10 deficiency increases mortality in male C57BL/6 mice after SARS-CoV-2 infection. Mice were intranasally infected with 8.62 × 10^4^ TCID_50_ of MA10 SARS-CoV-2. **(A-D)** Mortality. Mice were monitored over 12 days for survival. **(E-H)** Body weight. Percentage of body weight of infected mice were recorded. The genotypes of the mice are given on the right side of the respective graphs. Non-S: Non-survivors and S: Survivors. Results are summarized from 3-7 independent experiments. Data in (E-H) are represented as mean ± SEM. Statistical significance was determined by Kaplan-Meier survival curve analysis. *, *P* < 0.05; ****, *P* < 0.0001; ns: not significant.

### Cxcl10 deficiency does not alter histopathological damage and SARS-CoV-2 load in mouse lung

To determine the effect of *Cxcl10* deficiency on histopathology, inflammation and virus load after MA10 SARS-CoV-2-infection, we harvested lung tissues on day 4 post infection, the day before the first death occurs in the model. In both *Cxcl10*^+/+^ and *Cxcl10*^*-*/-^ mice, the most noticeable microscopic changes in lung were observed in the medial aspect (central hilum region) and to a lesser extent at the periphery of the lobe, which consisted of bronchointerstitial pneumonia characterized by increased alveolar thickening compared to uninfected lung (Fig. 2A-D). We also observed peribronchiolar and perivascular edema with infiltrating inflammatory cells and cellular debris (inflammatory exudate in multiple large and smaller bronchioles) (Fig. 2A-D). Based on various parameters, cumulative histopathological scores were assigned to the lung samples (see S1 Table). Overall, the microscopic changes in the lung were similar between infected *Cxcl10*^+/+^ and *Cxcl10*^*-*/-^ mice at this timepoint (Fig. 2E) and consistent with those previously described in the mouse adapted strain of SARS-CoV-2. We also performed histopathological evaluation of kidney, heart, brain and spleen from these mice. No significant lesions attributable to SARS-CoV-2 infection were present in the organ sections (data not shown). Notably, in the kidneys of both *Cxcl10*^+/+^ and *Cxcl10*^*-*/-^ mice, cytoplasmic vacuolization of cortical tubule epithelial cells throughout the renal cortices was a consistent yet unsuspected finding. The renal cortical tubular cytoplasmic vacuolation was deemed a background lesion unrelated to experimental SARS-CoV-2 infection since uninfected *Cxcl10*^+/+^ and *Cxcl10*^*-*/-^ control mice had similar renal changes. The levels of *Il6* and *Ifng* transcripts were also similar in the two groups, whereas *Tnfα* mRNA levels were ∼2-3-fold higher in *Cxcl10*^*-*/-^ mice compared to *Cxcl10*^+/+^ controls (Fig. 2F). Finally, we quantified the lung viral load by qPCR and found similar levels of SARS-CoV-2 N1 gene expression in infected *Cxcl10*^+/+^ and *Cxcl10*^*-*/-^ mice at 4 days post infection (Fig. 2F).

**Figure 2.**
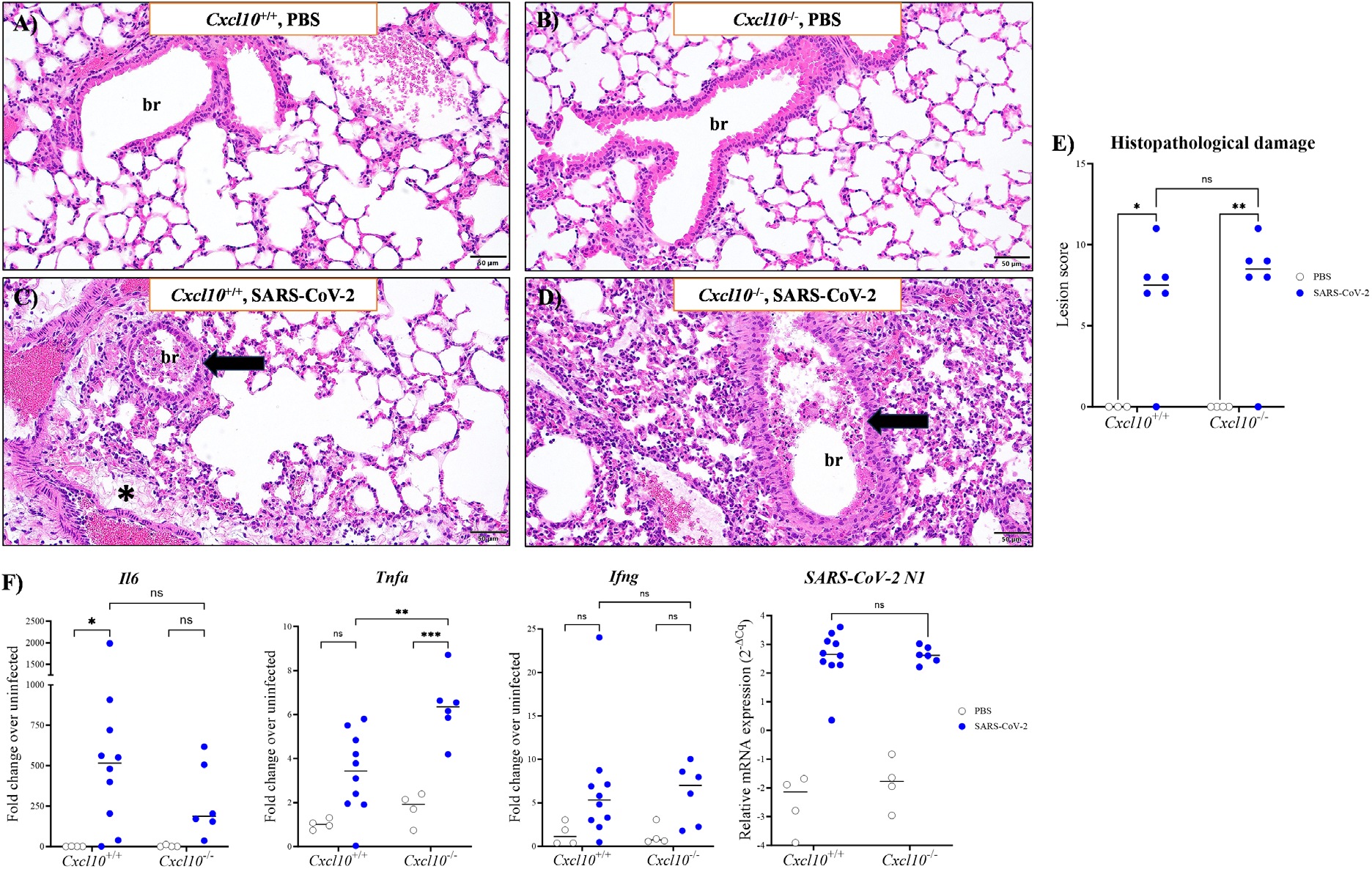
Effect of Cxcl10 deficiency on lung histopathology, inflammatory cytokine expression and SARS-CoV-2 viral load in infected mice. Male *Cxcl10*^*+*/+^ and *Cxcl10*^*-*/-^ mice were intranasally infected with SARS-CoV-2 or an equal volume of PBS and analyzed 4 days post infection. **(A-D)** H&E at 20x magnification. Representative sections of lung tissue are shown. The genotype and challenge are indicated at the top of each panel. Peribronchiolar and perivascular edema with infiltrating inflammatory cells are indicated with an asterisk; br: bronchiole. Arrows indicate cellular exudate that partially occludes a bronchiole lumen. **(E)** Pathology score of the lung tissues assigned in a blinded manner (see Table S1 for detailed score assignment). **(F)** Gene expression in the lungs of PBS and SARS-CoV-2 inoculated mice (legend) analyzed by qPCR for the indicated genes. Results are a summary of data from 4 independent experiments with 3-10 mice per group. Each data point in panel F is derived from an individual mouse and the median value is indicated by a black horizontal line. The infection status of the mice is given on the right side of the figure. Statistical significance was calculated using two-way ANOVA. *, *P* < 0.05; **, *P* < 0.01, ***, *P* < 0.001, ****, *P* < 0.0001; ns: not significant.

### Reduced B and T lymphocyte accumulation in SARS-CoV-2-infected lung from Cxcl10-deficient mice

CXCR3 and its ligands CXCL9, CXCL10 and CXCL11 are known to regulate leukocyte recruitment during numerous viral infections ^14^. On day 4 post infection, we found that in our model, *Cxcr3* transcript levels were decreased in both infected *Cxcl10*^*+/+*^ and *Cxcl10*^*-*/-^ mouse lung relative to the uninfected controls (Fig. 3A). In contrast, expression of all 3 chemokine ligands of Cxcr3 was strongly increased in infected *Cxcl10*^*+/+*^ mice compared to the PBS controls. Interestingly, while lung *Cxcl11* expression was comparable in infected *Cxcl10*^*+/+*^ and *Cxcl10*^*-*/-^ mice, the latter displayed a significantly higher induction of *Cxcl9* than infected *Cxcl10*^*+/+*^ mice (Fig. 3B-D).

**Figure 3.**
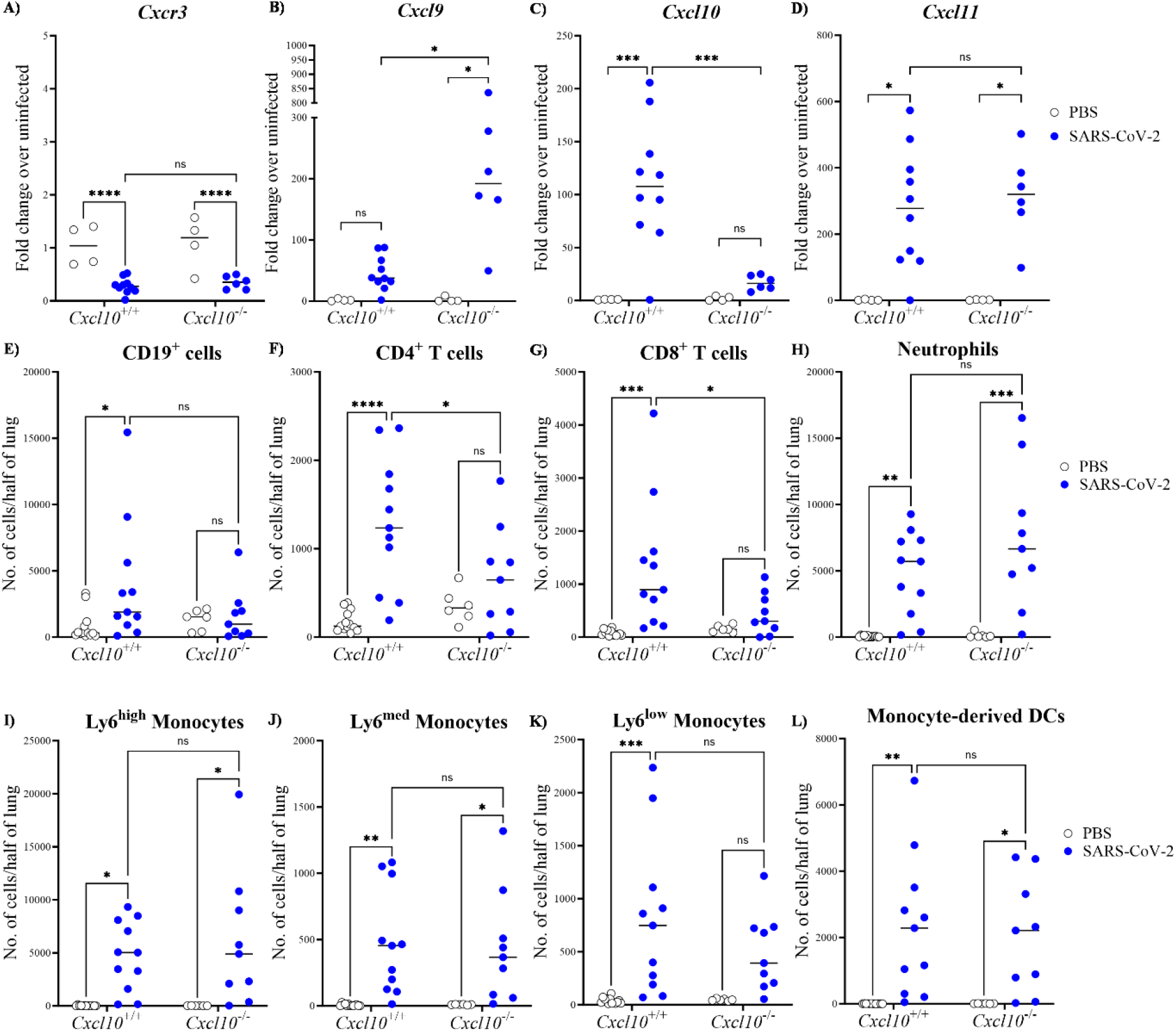
Lymphocyte accumulation is impaired in SARS-CoV-2 infected *Cxcl10*^-/-^ *mouse lung*. Male *Cxcl10*^+/+^ and *Cxcl10*^-/-^ mice were intranasally infected with SARS-CoV-2 or an equal volume of PBS, as indicated in the legends on the right side of each graph row. **(A-D)** Gene expression analysis in the lung at 4 days post infection by qPCR of the genes indicated above each graph. **(E-L)** Lung parenchymal leukocyte composition. Mice were intravenously injected with an anti-CD45-BV421 antibody for intravital staining 5 minutes prior to euthanization. Lung samples were collected, enzymatically digested and stained to quantify content of the leukocyte subtypes indicated at the top of each panel. Results in panels A to L are summary data from at least 4 independent experiments with 4-11 total mice per group, where each data point is derived from an individual mouse and the median value is indicated by a black horizontal line. The infection status of the mice is given on the right side of the figure. Statistical significance was calculated using two-way ANOVA. *, *P* < 0.05; **, *P* < 0.01, ***, *P* < 0.001, ****, *P* < 0.0001; ns: not significant.

Next, we studied leukocyte accumulation in the lung. To distinguish circulating from parenchymal cells within the lung, we performed intravital staining with an anti-CD45 antibody ^15^ (see Fig. S1 for gating strategy). The absolute numbers of parenchymal CD19^+^ B cells and both CD4^+^ and CD8^+^ T lymphocytes were increased in *Cxcl10*^+/+^ mice at day 4 post infection compared to uninfected control mice. In contrast, no significant lymphocyte recruitment occurred in infected *Cxcl10*^-/-^ mice compared with their uninfected littermate controls (Fig. 3E-G). Consistent with previous reports, the absolute numbers of neutrophils, monocytes and monocyte-derived dendritic cells increased within *Cxcl10*^+/+^ lung parenchyma post infection ^15^; however, we found no difference in their accumulation between infected *Cxcl10*^+/+^ and *Cxcl10*^-/-^ mice (Fig. 3H-L). Therefore, Cxcl10 appears to promote lymphocyte recruitment in the mouse lung after SARS-CoV-2 infection.

## Discussion

Due to the repeated emergence of variants and subvariants of SARS-CoV-2 and the waning immunity from past infections or vaccinations in the population, COVID-19 has remained endemic worldwide. Therefore, continued efforts to thoroughly investigate and validate potential therapeutic targets is important to advance diverse tools to reduce disease burden. Circulating CXCL10 has been positively correlated with the development of COVID-19 ARDS, suggesting the possible value of this chemokine as a biomarker of disease and therapeutic target ^3^. Contrary to this notion, our findings demonstrate that Cxcl10 sufficiency may actually be protective in reducing SARS-CoV-2-induced mortality in mice independent of lung damage, inflammation, and virus burden. Thus, our preclinical data in our model do not support targeting CXCL10 in severe COVID-19.

While we have not established the precise mechanism by which Cxcl10 may be protective in the model, we found that Cxcl10 deficiency blunted or ablated the normal lymphocyte infiltrate in the lung parenchyma after SARS-CoV-2 infection. This is consistent with the expression of the Cxcl10 receptor, Cxcr3, on these cells in a typical antiviral Th1 skewed immune response. Interestingly, we found that *Cxcl9*, another Cxcr3 agonist, was markedly upregulated in infected *Cxcl10*^-/-^ mice compared to infected *Cxcl10*^+/+^ control mice; however, this was not sufficient to compensate for the lack of Cxcl10 to recruit lymphocytes or protect against increased mortality. Although we did not detect a Cxcl10-dependent difference in viral burden at day 4 post infection, it is important to note that we only quantified the viral N gene by qPCR and not replication competent virus. Moreover, the effects on viral burden from defective lymphocyte accumulation in the lung might not be apparent until later time points when adaptive immunity occurs and mortality differences become more obvious. In this regard, a coordinated adaptive immune response is known to limit the severity of COVID-19 ^16,17^, and lymphocyte-deficient SCID and *Rag*^-/-^ mice are unable to clear SARS-CoV-2 infection ^18,19^.

In our model, the harmful effect of Cxcl10 deficiency on mortality was found for male but not female mice. In humans, PBMCs from healthy young male donors have been reported to secrete higher amounts of CXCL10 when compared to PBMCs from females upon stimulation with irradiated SARS-CoV-2 for 24 h but not after 7 days, which suggests that the kinetics of CXCL10 induction may vary between the two sexes ^20^. Likewise, it is possible that the dynamics of Cxcl10 expression may vary between male and female SARS-CoV-2 infected mice, affecting disease morbidity and mortality. Additionally, consistent with the increased COVID-19 age-adjusted mortality in men, females may possess more redundant mechanisms for host defense against SARS-CoV-2. We can speculate that an initial short-lived CXCL10 response may be beneficial whereas persistently sustained CXCL10 amounts, as seen in COVID-19 patients, may contribute to dysregulated inflammation and ARDS development.

In addition to its role in the regulation of lymphocyte migration, CXCL10 might exert some receptor-independent effects that could explain the differences reported here. For instance, CXCL10 is known to compete with cell surface heparan sulfate to inhibit dengue virus binding to the Hepa1-6 mouse hepatoma cell line ^21^, and the virus load is significantly higher in the brain of *Cxcl10*^-/-^ mice ^22^. However, CXCL10 is unlikely to exert direct antiviral effects on SARS-CoV-2 since in our study the lack of Cxcl10 did not increase the viral load in mouse lung tissue. Consistent with our results, previous studies have reported no correlation between alveolar concentrations of SARS-CoV-2 and alveolar CXCL10 amounts ^3^. In addition, CXCL10 upregulation in Calu-3 human lung epithelial cells is regulated by the AKT pathway. Application of an AKT inhibitor reduced CXCL10 induction but did not lower the viral RNA load ^23^.

Our study has important limitations. First, the mouse model we used may have adapted to constitutive and complete Cxcl10 deficiency. Future work identifying the precise cell types responsible for Cxcl10 production will enable follow-up studies targeting Cxcl10 in these cells using conditional knockout mice. Second, our study is limited to events at baseline and day 4 post infection, the day before most infected mice die. While it is difficult to control such studies once mortality begins because of survivor bias, there may be important Cxcl10-dependent immunological events occurring at later timepoints that are key to determining the precise survival outcome. Third, we have not quantified all the leukocyte subsets that may be responsive to Cxcl10. For instance, CXCL10 regulates chemotaxis of NK cells ^24^. Quantification of NK cells in *Cxcl10*^-/-^ mice would be important for future work since NK lymphopenia and dysfunction are observed during COVID-19 ^25^. T cells are required for SARS-CoV-2 virus clearance in mice ^26^ and therefore studying the SARS-CoV-2-specific adaptive immune response in the *Cxcl10*^-/-^ mice will be of interest. However, since Cxcr3 is expressed on multiple cell types, it is beyond the scope of the present study to target each one for their contribution to survival outcome. Fourth, we face the common difficulty in survival studies of determining the precise cause of death. While we have focused our attention on the lung, there may be important pathological events occurring in other organs that we missed. Nevertheless, we did not observe immunopathology in any of the other organs we examined (heart, kidney, brain and spleen). Fifth, we have studied only young mice. The importance of Cxcl10 may differ in older mice and in humans of different ages and may be influenced by the presence of comorbidities.

In conclusion, our study suggests that Cxcl10 mediates pan-lymphocyte accumulation in lung and may be protective against fatal outcome after SARS-CoV-2 infection in mice. If translated to the more complex human setting, these preclinical results would not support targeting CXCL10/CXCR3 axis in severe COVID-19. Considering both the available human and mouse data, CXCL10 likely plays a complex role during development of a dysregulated immune response and virus clearance by mediating lymphocyte trafficking within lung, causing an interplay between both processes, thus influencing disease outcome. Future studies will be necessary to dissect mechanisms that confer CXCL10-dependent host protection during early and late phases of infection.

## Materials and methods

### Mice

Mice aged 4-6 months were used. WT C57BL/6 (JAX664, *Cxcl10*^+/+^*Cxcl11*^-/-^), *Cxcl10*^-/-^ (B6.129S4-*Cxcl10*^*tm1Adl*^/J, *Cxcl10*^-/-^*Cxcl11*^+/+^) and 129S1/SvImJ (*Cxcl10*^+/+^*Cxcl11*^+/+^) mice were purchased from The Jackson Laboratory (Bar Harbor, ME, USA). To obtain *Cxcl10*^+/+^*Cxcl11*^+/+^ mice on the JAX664 genetic background, 129S4/SvLaeJ and C57BL/6 mice were crossed to select for *Cxcl11*^+/+^ offspring, which were backcrossed to C57BL/6 mice for 8 generations. The following primers were used to genotype the mutant and the WT *Cxcl11* genes in C57BL/6 and 129S4/SvLaeJ mice, respectively: C57BL/6 Forward, 5’-AGCCATAGCCCTGGCTGCAATA-3’; 129S4/SvLaeJ Forward, 5’-CCATAGCCCTGGCTGCGATC-3’; and common reverse, 5’-TGCTGCGCCATTGCTAGAACTTC-3’. The mice were maintained and bred under specific pathogen-free conditions with 12-hour light/dark cycle and access to food and water *ad libitum*. The studies were performed in accordance with NIH institutional guidelines and were conducted in Assessment and Accreditation for Laboratory Animal Care-accredited Biosafety Level (BSL) 2 and 3 facilities at the NIAID/NIH using a protocol approved by the NIAID Animal Care and Use Committee.

### Virus stock preparation and mouse infections

Virus stock generation and mouse infections were performed in BSL-3 containment laboratories. The MA10 strain of SARS-CoV-2 was obtained from the Biodefense and Emerging Infections Research Repository (BEI resources, Manassas, VA, USA). The virus stocks were generated by infections in Vero E6 cells expressing TMPRSS2 ^27^ at a multiplicity of infection of 0.01, at 37ºC for 48 h in 5% CO_2_. Media were harvested and centrifuged to remove cellular debris and collect the virus. Sequencing was performed to confirm single nucleotide variants consistent with MA SARS-CoV-2.

For intranasal infections, mice were anesthetized by isoflurane inhalation. A dose of 8.62 × 10^4^ TCID_50_ of MA10 SARS-CoV-2 was administered to mice. Mock-infected mice received the equivalent volume (20 µL) of sterile PBS intranasally. Mortality, pain score and body weights of mice were monitored throughout the experiment, and mice demonstrating a pain score of 3 (shaking/shivering, eyes closed/shut, labored breathing and reluctance to move; unable to reach for sustenance) and/or body weight drop by ≥ 25% of the initial body weight were euthanized per protocol.

### Chemokine, cytokine and viral load analysis by qPCR

The inferior and post-caval lobes of lungs were collected for gene expression analysis. Samples were homogenized in 1 mL of HBSS (Thermo Fisher Scientific, Waltham, MA, USA) using 2 mL tubes containing zirconium-silicate spheres (MP Biomedicals, Irvine, CA, USA) and Bead Ruptor 24 (OMNI International, Kennesaw, GA, USA) at 5.15 m/s, 15 s, 1 cycle. For virus inactivation, 250 µL of the tissue homogenate were added to 750 µL of TRIzol LS (Thermo Fisher Scientific) and kept at room temperature for 10 mins. Post chloroform extraction, the aqueous phase was mixed 1:1 v/v with 70% ethanol and loaded into an Isolate II RNA Mini kit column (Bioline, Cincinnati, OH, USA). Total RNA was purified according to the manufacturer’s instructions. cDNA synthesis was performed using the SensiFAST cDNA Synthesis kit (Bioline) and 200 ng of RNA as template, and qPCR reactions were run in a CFX96 Touch Real-Time PCR instrument (Bio-Rad, Hercules, CA, USA) using the SensiFAST SYBR Hi-ROX kit (Bioline) and predesigned KiCqStart SYBR green primers for the genes of interest (MilliporeSigma, St. Louis, MO, USA). The expression fold change over corresponding mock-infected group for every gene was calculated as 2^-(ΔΔCt)^ using the geometric mean Ct of 3 genes, *Gapdh, ActB* and *Hprt1* as reference. Viral load quantification was conducted using Primetime One-step RT-qPCR Master Mix (Integrated DNA Technologies, Coralville, IA, USA). Quantification of the nucleocapsid (N) gene of SARS-CoV-2 was performed using the 2019-nCoV RUO Kit (Catalog# 10006713, Integrated DNA Technologies). The primers for *Gapdh* (Mm01180221_g1) were purchased from Thermo Fisher Scientific.

### Histopathology

Under BSL-3 conditions, superior and middle lobe of lung, whole spleen, kidney, brain and heart were collected and immersed and fixed in 10% neutral buffered formalin (MilliporeSigma) for a minimum of 3 days for virus inactivation. Subsequently, the tissues were transferred into 70% ethanol and were dehydrated through graded alcohols, cleared in xylene, and then infiltrated with paraffin. All tissues were embedded in paraffin after processing. The paraffin blocks were cut into 5 µm sections using a microtome. The unstained slides were then deparaffinized using xylene and graded alcohols to water and stained in hematoxylin, and rinsed again in water. Next, the slides were placed in 95% ethanol prior to staining with eosin and dehydrated through graded alcohols to xylene. Finally, the stained H&E slides were mounted with permount. Blinded tissue sections were evaluated by a board-certified veterinary pathologist (DAA) using an Olympus BX51 light microscope, and photomicrographs were taken by an Olympus DP28 camera.

### Flow cytometry

To selectively stain leukocytes in circulation, mice were anesthetized with isoflurane, and CD45-BV421 antibody (100 µL/mouse, 1:20 dilution in PBS) was injected retroorbitally. Mice were allowed to recover for 5 min and then euthanized. The left lung was collected and minced into small pieces prior to enzymatic digestion with 125 µg Collagenase D (MilliporeSigma) in Dulbecco’s Modified Eagle Medium (Thermo Fisher Scientific) at 37ºC for 30 min prior to passing the tissue through a 100 µm cell strainer by gently pressing with a piston into a 50 mL falcon tube containing 10 mL of FACS buffer (PBS supplemented with 2% FCS and 2 mM EDTA). The cell pellets were treated with ACK lysis buffer (Quality Biological Inc., Gaithersburg, MD, USA), washed and resuspended in FACS buffer. The cells were stained with a viability dye (LIVE/DEAD™ fixable aqua dead cell stain, Thermo Fisher Scientific), Fc blocker (Clone: 93, BioLegend, San Diego, CA, USA) and fluorescently-labeled antibodies with Super Bright complete staining buffer (Thermo Fisher Scientific) at room temperature for 30 min. The samples were washed with PBS and the cells were treated with 4% PFA (Thermo Fisher Scientific) for 30 min at room temperature for virus-inactivation. Finally, the cells were washed and resuspended in PBS. SPHERO™ AccuCount Particles (Spherotech, Inc., Lake Forest, IL, USA) were added to the samples during acquisition for cell quantification. BD Fortessa and FlowJo (BD Bioscience, Franklin Lakes, NJ, USA) were used for data acquisition and analysis, respectively. The following antibodies were used: CD4-PerCP/Cy5.5 (Clone: RM4-5), CD8a-PE/Cy7 (Clone: 53-6.7), CD19-FITC (Clone: 6D5), CD19-PE/Cy7 (Clone: 6D5), CD45-BV421 (Clone: 30-F11), CD103-PE/Cy7 (Clone: 2E7), Ly6C-PerCP/Cy5.5 (Clone: HK1.4), MHC-II-BV785 (Clone: M5/114.15.2), NK1.1-FITC (Clone: PK136), TCRβ-APC/Cy7 (Clone: H57-597), TCRβ-FITC (Clone: H57-597) and TCRγ/δ-FITC (Clone: GL3) were purchased from BioLegend, CD8a-BV786 (Clone: 53-6.7) was purchased from BD Biosciences, and CD11b-SB600 (Clone: M1/70), CD45-BV395 (Clone: 30-F11), CD170 (Siglec F)-BV711 (Clone: 1RNM44N) were purchased from Thermo Fisher Scientific.

### Statistical analysis

GraphPad Prism (GraphPad Software, La Jolla, CA, USA) was used to graph the data and calculate statistical differences. The statistical tests used are indicated in each figure legend.

## Acknowledgments

We thank the staff of the NIAID BSL-2 and BSL-3 facilities.

## Author Contributions

PMM conceptualized the study. SM, JDW, SMP, MM, XL and GH performed the experiments. RJ and HZ provided the resources. SM, JDW, SMP, J-LG, BLK, JMF, DAA and PMM analyzed the data. All authors have read and agreed to the final version of the manuscript.

