## Supplementary material for "Cxcl10 is required for survival during SARS-CoV-2 infection in mice": Fig. S1

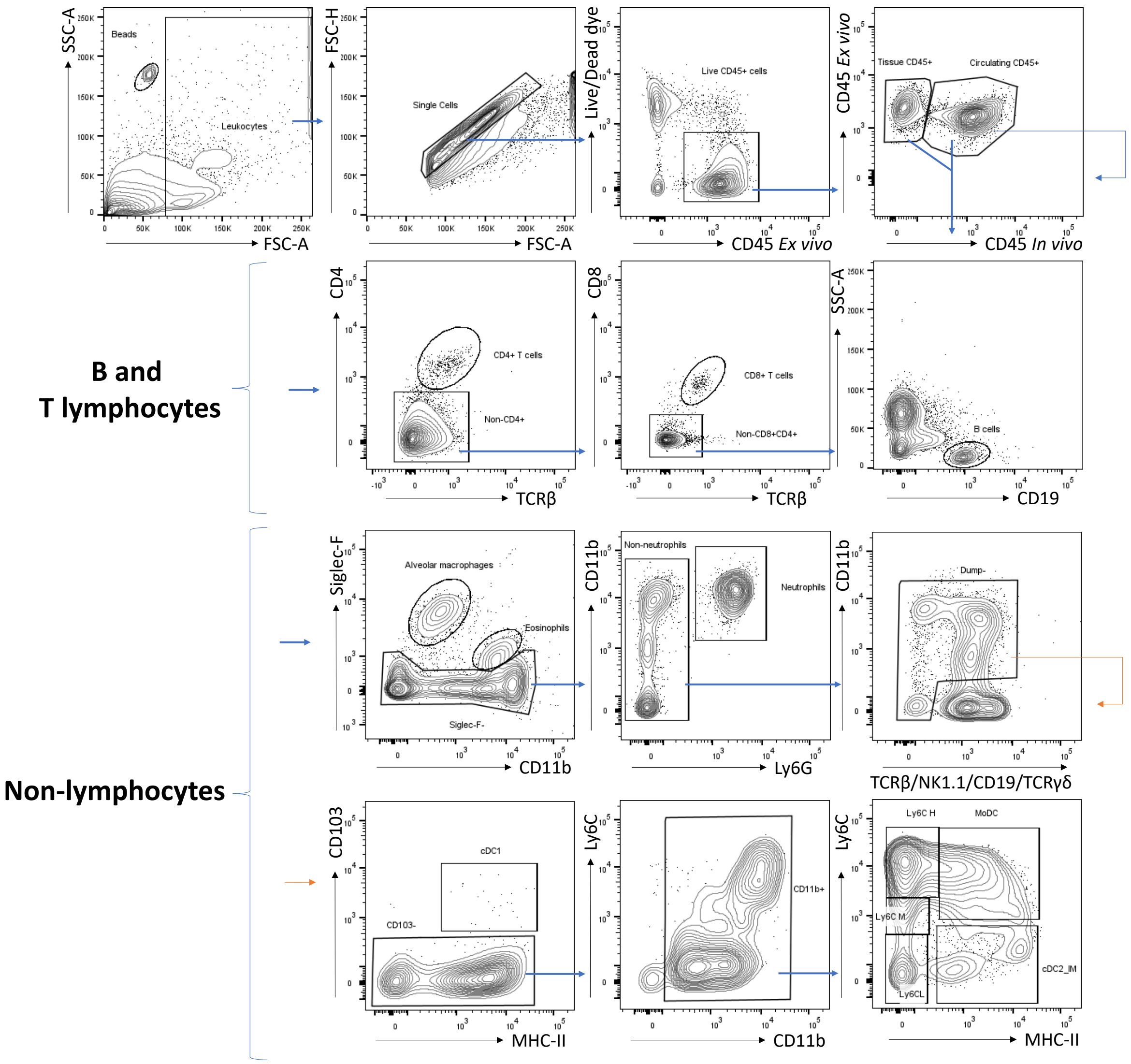


**Fig S1: Gating strategy for analysis of leukocytes in mouse lung.** On side scatter-area (SSC-A) and forward scatter-area (FSC-A) plots, the counting beads were gated to obtain absolute count of leukocytes for analysis. The total leukocytes acquired were gated and then the singlets were identified on the forward scatter-height (FSC-H) vs. FSC-A plots. Next, the live, total $CD45^+$ cells were selected. Lung parenchymal cells ($CD45^+$ *ex vivo*) were differentiated from circulating leukocytes ($CD45^+$ *in vivo*). The subsequent gates were made for both parenchymal and circulating cells. In the lymphocyte panel, $CD4^+$ T cells ($CD4^+TCR\beta^+$) were selected, while the remaining events were gated to select for the $CD8^+$ T cells ($CD8^+TCR\beta^+$), and then the B cells ($CD19^+$). In the non-lymphocyte panel, alveolar macrophages were identified as Siglec-$F^+CD11b^{low}$ and the eosinophils as Siglec-$F^+CD11b^+$. The remaining events were gated to identify neutrophils ($CD11b^+Ly6G^+$). The non-lymphocytes and non-neutrophils were selected for downstream analysis ($CD11b^+TCR\beta^-TCR\gamma/\delta^-CD19^-NK1.1^-$). The conventional dendritic cells 1 (cDC1) were identified as $CD103^+$MHC-$II^+$, while the remaining cells expressing CD11b were gated to quantify $Ly6C^{low}$, $Ly6C^{medium}$ and $Ly6C^{high}$ monocytes, monocyte-derived dendritic cells (moDC) and conventional dendritic cells 2 or inflammatory macrophages (cDC1-IM).
